# Do sun orchids mimic buzz-pollinated plants? An experimental test of the adaptive significance of false anthers

**DOI:** 10.1101/2025.03.15.643429

**Authors:** Daniela Scaccabarozzi, Nina Sletvold

**Author notes:** For correspondence:* Uppsala University, Department of Ecology and Genetics Evolutionary Biology Centre Address: Norbyvägen 18 D, 752 36 Uppsala.

## Abstract

1. Mimicry implies that an organism gains fitness by resembling a model species, and one example is rewardless plants that attract pollinators by resembling co-flowering species that provide rewards. While trait matching between mimic and model has been characterised in many cases of putative floral mimicry, few have demonstrated that resemblance is adaptive and dependent on model presence.
2. Sun orchids (*Thelymitra*) are believed to mimic flowers of buzz-pollinated rewarding plants by displaying false anthers. To test the adaptive value of the false anthers we examined whether fruit production of *T. crinita* and *T. macrophylla* was reduced when anthers were experimentally removed or obscured, and whether the reduction was stronger when putative model plants were abundant. We also assessed visual flower similarity of both orchids and their putative model plants according to bee colour perception and identified shared pollinators and whether their behaviour on *T. crinita* was similar to that on co-flowering rewarding plants.
3. Fruit production of both sun orchids was strongly reduced (60-71%) by removal or painting of false anthers but was not affected by the abundance of model plants. Sun orchid flowers closely matched flower colour of co-flowering pollen-rewarding species, and *T. crinita* shared pollinators with the rewarding species. Visiting bees attempted to buzz and manipulate the false anther, with a behaviour similar to that observed on model plants.
4. The experimental results demonstrate that the false anther is an important adaptation to pollination in sun orchids. Striking visual flower similarity and shared pollinators between orchids and models suggest that sun orchids are pollinated by bees that mistake orchids for buzz-pollinated rewarding plants. The adaptive value of the false anther did not depend on model plant abundance in the local population, indicating that the relevant spatial scale is larger, or that the effects of the model species are weak in comparison to effects of other rewarding species, i.e., that magnet effects of nectar-rewarding species are dominating.
5. False anthers are widespread in the genus *Thelymitra*, and this “mimicry trait” seems to represent an evolutionary novelty that offers unique opportunities to explore adaptations to pollination in deceptive plants.

## 1 INTRODUCTION

Mimicry provides a multitude of fascinating examples of adaptation by natural selection, spanning the tree of life (Joron & Mallet, 1998; Dalziell & Welbergen, 2016; Jamie, 2017; de Jager & Anderson, 2019). In plants, some of the most well-known cases include rewardless plants that attract pollinating insects by resembling co-flowering rewarding species (Renner, 2006; Johnson & Schiestl, 2016). This is known as Batesian floral mimicry and occurs in 32 plant families (Renner, 2006). Floral mimicry may range from a general resemblance of rewarding species to precise trait matching of a specific model species, i.e., from ‘generalized food deception’ to true Batesian mimicry (Johnson & Schiestl, 2016). Common to all cases is that mimetic resemblance should be adaptive. In addition, the mimic’s fitness benefit of similarity is expected to be highest when the model is abundant relative to the mimic, because pollinator mistakes should be frequent in such conditions (Johnson, 1994; 2000).

Despite a sustained interest in floral mimicry, few studies have demonstrated its adaptive significance or context-dependence. Many studies conclude on mimicry based on documentation of overlap between a putative model and mimic in floral traits and phenology, in combination with pollinator sharing (Jersáková et al., 2009; Scaccabarozzi et al., 2018; 2020). We thus lack a clear understanding of the floral traits involved, and of how model abundance affects the adaptive value of traits. To conclusively link fitness variation in the mimic to degree of trait matching of a putative model, manipulations of key phenotypic traits is a powerful tool. This approach is rare, but notable exceptions include manipulations of inflorescence shape (Johnson et al., 2003), flower colour (Jersáková et al., 2012), and colour patterns (Scaccabarozzi et al., 2023) in mimics. Similarly, a few studies altering model abundance (Anderson & Johnson, 2006) or distance to mimic (Peter & Johnson, 2009; Duffy & Johnson, 2017) have found fitness of the mimic to increase with model abundance or proximity. However, it is generally unknown if this is mediated by trait similarity of the model species, or by increased pollinator attraction, caused by a ‘magnet effect’ (Thomson, 1978; Feinsinger, 1987; Laverty, 1992; Johnson et al., 2003a). This could be clarified by trait manipulations in different rewarding contexts.

Mimics probably converge on multiple visual signals of the model, and suggested cases of floral mimicry range from resemblance of the entire inflorescence and flower structure, shape and colour (Johnson et al., 2003b; Jersáková et al., 2016), to the imitation of flower components, such as pollen-like papillae, false anthers, and false nectar guides (Lunau et al., 2020; 2021; 2024). In bee-pollinated species, stamen-like structures associated with pollen as a reward can constitute an important visual signal, and studies using false colour photography to identify colour patterns according to the bee view have suggested stamen mimicry to be the widest distributed mimicry system worldwide (Lunau et al., 2021; 2024). In most cases, stamen mimicry involves trait convergence in rewarding species (i.e., Müllerian mimicry), and while the functional (Walker-Larsen & Harder, 2001; Dieringer & Cabrera, 2002) and adaptive (Newman et al., 2022; Wen et al., 2024) roles of staminoides have been experimentally demonstrated in a few nectar-producing species, this is largely unexplored in deceptive systems.

Some of the most striking potential cases of stamen mimicry occur in the global biodiversity hotspot of the Southwestern Australia floristic region (SWAFR) (Duncan et al., 2004; Houston, 2018; Scaccabarozzi et al., 2018; 2020; Lunau et al., 2021). In this region, many deceptive species have bright yellow structures that resemble anthers of buzz-pollinated flowers (Bernhardt and Burns-Balogh, 1986), i.e., pollen-rewarding flowers that require the production of intense bee vibrations to release pollen (De Luca & Vallejo-Marín, 2013). Buzz-pollinated flowers are characterised by poricidal, conical anthers, that often form part of the attractive floral display (Buchmann, 1983; Vallejo-Marín, 2019), and similar structures are for example found in the deceptive genus *Thelymitra* (sun orchids, Orchidaceae). Bee-pollinated species in this genus have been suggested to mimic anthers of several co-blooming and buzz-pollinated rewarding flowers, including resemblance between *Thelymitra nuda* and the genera *Dichopogon* and *Thysanotus* (Asparagaceae) (Bernhardt & Burns-Balogh, 1986), and between *T. macrophylla* and *Orthrosanthus laxus* (Iridaceae) (Edens-Meier et al., 2014). However, whether bee visitors engage in pollen-gathering behaviour, and whether false anthers are crucial for sun orchid pollination success, is presently unknown.

The aim of this study was to experimentally test whether false anthers in *Thelymitra* orchids represent buzz-pollination mimicry. To achieve this, we first assess visual similarity of flowers of *Thelymitra* and putative model plants according to bee colour perception and identify shared pollinators and assess their behaviour on orchids and model plants. Second, we manipulate the display of false anthers in *Thelymitra* in two rewarding contexts, to specifically test whether (i) orchid fitness is reduced when anthers are removed or obscured, and (ii) the reduction is stronger when putative model plants are abundant.

## 2 METHODS

### 2.1 Study system

*Thelymitra* is a genus of over 100 nectarless orchid species primarily found in Australia. These terrestrial, perennial herbs have inconspicuous roots, oval-shaped tubers, and a single leaf near the base of the plant (Western Australian Herbarium, 1998). Most species within the genus are believed to be bee-pollinated via food deception, although some may be autogamous (Beardsell & Bernhardt, 1983; Edens-Meier et al., 2014). Unlike typical orchids, *Thelymitra*’s labellum resembles the other petals, and flowers have an almost radially symmetric perianth. The flower’s sexual parts are fused to a short column ending with a hood-like structure (mitra) or pseudanthery (Pridgeon et al., 2001; Fig. 1), hereafter ‘false anther’. Sun orchids, as their common name suggests, typically open their flowers in response to daylight, warm temperatures, and humidity. Here, we focus on two common species, *Thelymitra macrophylla* and *T. crinita*. The two species have similar flower morphology (Fig. 1), but *T. macrophylla* flowers earlier than *T. crinita*. In a previous study, bees of the genera *Leioproctus* and *Lasioglossum* were identified as pollinators of *T. macrophylla*, while only *Leioproctus* bees pollinated *T. crinita* (Edens-Meier et al., 2013).

**Figure 1.**
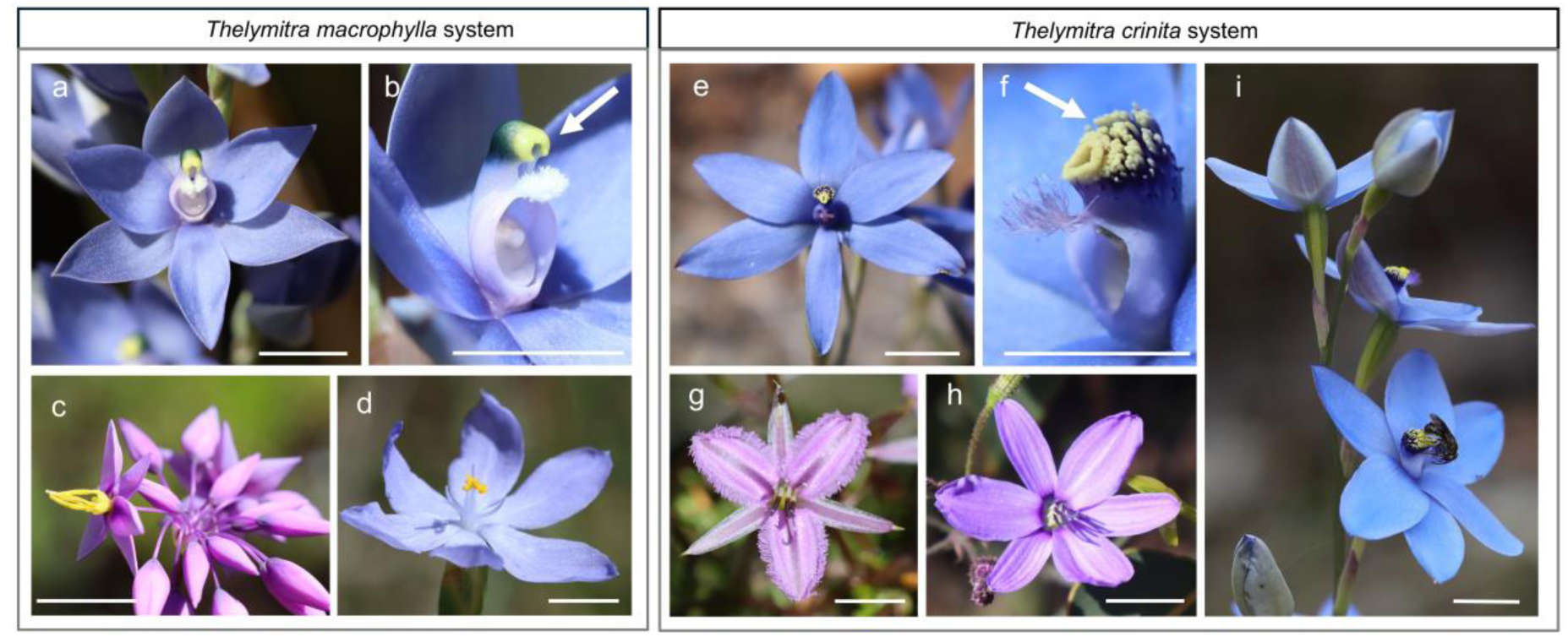
The sun orchids *Thelymitra macrophylla* (left panel) and *T. crinita* (right panel) with their main co-flowering, pollen-rewarding species. a: *Thelymitra macrophylla*; b: close-up of reproductive structures with false anther (arrow); c: *Sowerbaea laxiflora*; d: *Orthrosanthus laxus*; e: *Thelymitra crinita*; f: close-up of reproductive structures with false anther (arrow); g: *Thysanotus manglesianus*; h: *Agrostocrinum hirsutum*; i: native bee *Leioproctus* (Colletidae) facing downwards on a *T. crinita* flower while manipulating the false anther. Scale bar: 1 cm. Photo Credits: Daniela Scaccabarozzi (a-h) and Mick Hurdus (i).

### 2.2 Study sites

We collected data from September to November 2023 in Southwestern Australia. The distribution of *T. macrophylla* and *T. crinita* ranges from Perth to Albany, and from Gingin to Esperance, respectively. Study populations of *T. macrophylla* were in *Banksia* woodland habitats within Warwick and Shanton Park bushland reserves, and study populations of *T. crinita* were found in forests dominated by *Eucalyptus marginata* in the Darling Range on the Perth hills (Fig. S1). Due to an unusual heatwave that caused flowering of *T. crinita* in the Perth region to end early, we also included a population in the Margaret River region, at Flat Rock bushland, Augusta (Fig. S1). Both *Thelymitra* species are abundant in the study regions, with typical population sizes ranging from 20 to 100 plants. The flowering time ranges from September to October for *T. marcrophylla* and from October to November for *T. crinita* (Western Australian Herbarium, 1998). Both orchids co-occur with a range of buzz-pollinated rewarding species with similar flower colour (Edens-Meier et al., 2014; Brundrett et al, 2024). In the *T. macrophylla* study populations, main co-flowering species included *Orthrosanthus laxus* (Iridaceae)*, Sowerbaea laxiflora* (Asparagaceae), and *Thysanotus manglesianus* (Asparagaceae), that all are pollen-rewarding (Fig. 1). In the *T. crinita* populations, the main co-flowering species were pollen-rewarding *T. manglesianus* and *Agrostocrinum hirsutum* (Asphodelaceae) (Fig. 1), and nectar-rewarding *Lechenaultia biloba*, *Dampiera linearis*, and *Scaveola calliptera* (Goodeniaceae).

### 2.3 Flower colour

To characterise flower colour in the two deceptive orchids and the co-flowering pollen- and nectar-rewarding species, we collected inflorescences in the field and brought them to the laboratory, where we measured six flowers per species using a field spectrometer (JAZ, Ocean Insight, Orlando, FL). On each flower, we took one measurement on the surface of the tepals or petals. To quantify colour similarity according to bee visual perception, we used the conventional bee vision model (colour hexagon space) based on photoreceptor spectral sensitivity of honeybees (Chittka, 1992). We quantified colour similarity by calculating the mean Euclidian distance between hexagon colour loci of *Thelymitra* and putative models. In addition, we used false colour photography as per Lunau et al. (2021), to identify colour patterns and details in the central area of the orchid flowers. This technique combines colour and UV photographs and is a robust method for obtaining detailed information on flower colour patterns under natural light conditions in the field (Verhoeven et al., 2018; Lunau et al., 2020). The technique splits colour photos into the three colour channels blue, green, and red, and by combining the green- and blue-channel photos with the UV photos, the resulting assemblage illustrates how bees perceive flower colour, i.e., including UV and excluding red.

### 2.4 Pollinator observations

To identify the pollinators of sun orchids, we assessed the approach of all visitors to the flowers (attraction to the flower without necessarily landing), their behaviour and potential landing, and the capability to remove and deposit orchid pollinia, combining direct pollinator observations with camera recordings. We initially included both orchid species, but due to very low insect activity in *T. macrophylla* populations, we focused on *T. crinita*. We conducted observations in two *T. crinita* populations in the Perth hills, and in one population in Flat Rock, Augusta, between 5 October and 1 November 2023, and between 15 October and 1 November 2024. Because natural visitation rate was very low, we created experimental arrays of *T. crinita* inflorescences close to rewarding species, to increase attractiveness (Scaccabarozzi et al., 2020). Arrays were formed by three cut orchid inflorescences placed in glass vials (one inflorescences per vial, each with 4–6 flowers) spaced 10 cm apart, and located ca 1 m from flowering individuals of the main rewarding species, *T. manglesianus* and *A. hirsutum*. In Perth hills, we conducted 80 and 20 ten-minute trials in the first and second population, respectively, and in the Augusta population, we conducted 200 ten-minute trials. This resulted in 50 hours pollinator observations in total. In each period, we also recorded insect behaviour using an EOS M video camera (Canon, Tokyo, Japan) for subsequent viewing in slow motion. Observations were done between 9 am and 3 pm (typical flower anthesis) during sunny or partly sunny days, when temperatures were above 22°C (measured 20 cm above the ground with a Smartsensor AR827, Sinosource Scientific Company, China).

To determine if *T. crinita* shares pollinators with putative model species, we also conducted pollinator observations and camera recordings of *T. manglesianus* and *A. hirsutum,* during some of the orchid arrays. For both species, we performed 21 ten-minute observation periods, yielding a total of 3 hours and 30 min observations per species. When possible, pollinators of both *T. crinita* and the putative model plants were identified in the field. In other cases, specimens were netted and later identified at the Museum of Western Australia.

### 2.5 False anther manipulations

To test the adaptive significance of false anthers, we used two experimental approaches. First, we examined the effect of removing the false anther in two *T. macrophylla* populations. Second, we examined the effect of obscuring the false anther using paint in two populations of each of the two species. In the first case, physical damage may influence bee behaviour on flowers. In the latter case, false anthers remained intact, and only the visual signal was removed.

To test if adaptive significance depends on rewarding context, we conducted two of the experiments at sites with varying abundance of rewarding species. In the anther-painting experiment on *T. macrophylla*, we could not identify any workable sites with variation in model abundance, and the experiment was conducted in two population that both had abundant model plants. In *T. macrophylla,* the anther-removal experiment was conducted in one population with putative model plants (*O. laxus, S. laxiflora, T. manglesianus*) and in one without. In *T. crinita*, we conducted the anther-painting experiment in two populations that differed quantitatively in local abundance of putative model plants. In both *T. crinita* populations, we quantified abundance of model plants by estimating the number of flowers per species in five randomly distributed 5 × 5 m plots per population. We counted all flowering stems and all flowers on ten stems per plant, and estimated total number of flowers by multiplying number of stems by mean number of flowers per stem. We included all rewarding species with blue-violet flower colour as potential model and magnet plants, i.e., *A. hirsutum, D. linearis, L. biloba*, *S. calliptera*, and *T. manglesianus*. The mean number of rewarding flowers per plot was 3.9 times higher in the population with high model abundance (mean ± SD = 87.6 ± 15.0) compared to the population with low model abundance (mean ± SD = 22.6 ± 12.4), and the ratio of mimics to models were approximately half in the former (0.45 and 0.89, respectively).

a. *Anther removal experiment*

On 24 September 2023, we randomly tagged 50 plants in each of two *T. macrophylla* populations separated by approximately 1 km within the Warwik Bushland Reserve, Perth. Putative model plants were present in one of the populations. In each population, we randomly assigned half of the plants to an anther removal treatment, while the other half were used as controls (n=25 per treatment). In the anther removal treatment, we carefully excised the false anthers (i.e., the upper part of the column, Fig. S2) from all open flowers using scissors. Most flowers were already open at the time of the experiment, and any remaining buds were removed. Prior to conducting manipulations, we recorded number of flowers per individual and examined if flowers had been visited. All flowers with pollinia removal or deposition were marked and excluded from the treatments (no difference between treatments; *P* > 0.05). In both populations, we recorded fruit production of all plants two weeks after removing false anthers.

To test if anther removal affected the ability to produce fruits, we supplementally hand-pollinated all flowers on 16 additional plants in another population in the Warwick bushland reserve, where eight had their false anther removed as described above, and eight were unmanipulated controls. We recorded fruit production two weeks later. Anther removal in *T. macrophylla* did not affect the ability to produce fruits (one-way ANOVA; F_1,12_ = 0.35, *P* = 0.56).

a. b) *Paint experiment*

On 4 October 2023, we randomly tagged 40 plants at flowering peak in each of two *T. macrophylla* populations (both with abundant model plants) in Shenton Bushland Reserve, Perth. In each population, we randomly assigned half of the plants to an anther paint treatment, while the other half were used as controls (n=20 per treatment). In the paint treatment, we obscured false anthers in all open flowers by applying a colour that matched the colour of the column (Eraldo Di Paolo, Acrylic Paint, Warm Blue; colour match examined by spectrometer measurements and false colour photography; Fig. 2A). To control for potential side-effects of painting, we also applied paint to the backside of the corolla of all flowers in the control treatment. On 17 October 2023, we randomly tagged 50 plants at flowering peak in each of two *T. crinita* populations (high vs. low abundance of model plants) on the Darling Scarp, Perth Hills. In each population, we randomly assigned half of the plants to an anther paint treatment, while the other half were used as controls (n=25 per treatment). Treatments were conducted as described for *T. macrophylla*. Prior to manipulations, we recorded number of flowers per individual and marked and excluded all flowers with pollinia removal or deposition for both orchids (no difference between treatments; both *P* > 0.05). In both experiments, we recorded fruit production of all plants two weeks after the anther manipulations.

**Figure 2.**
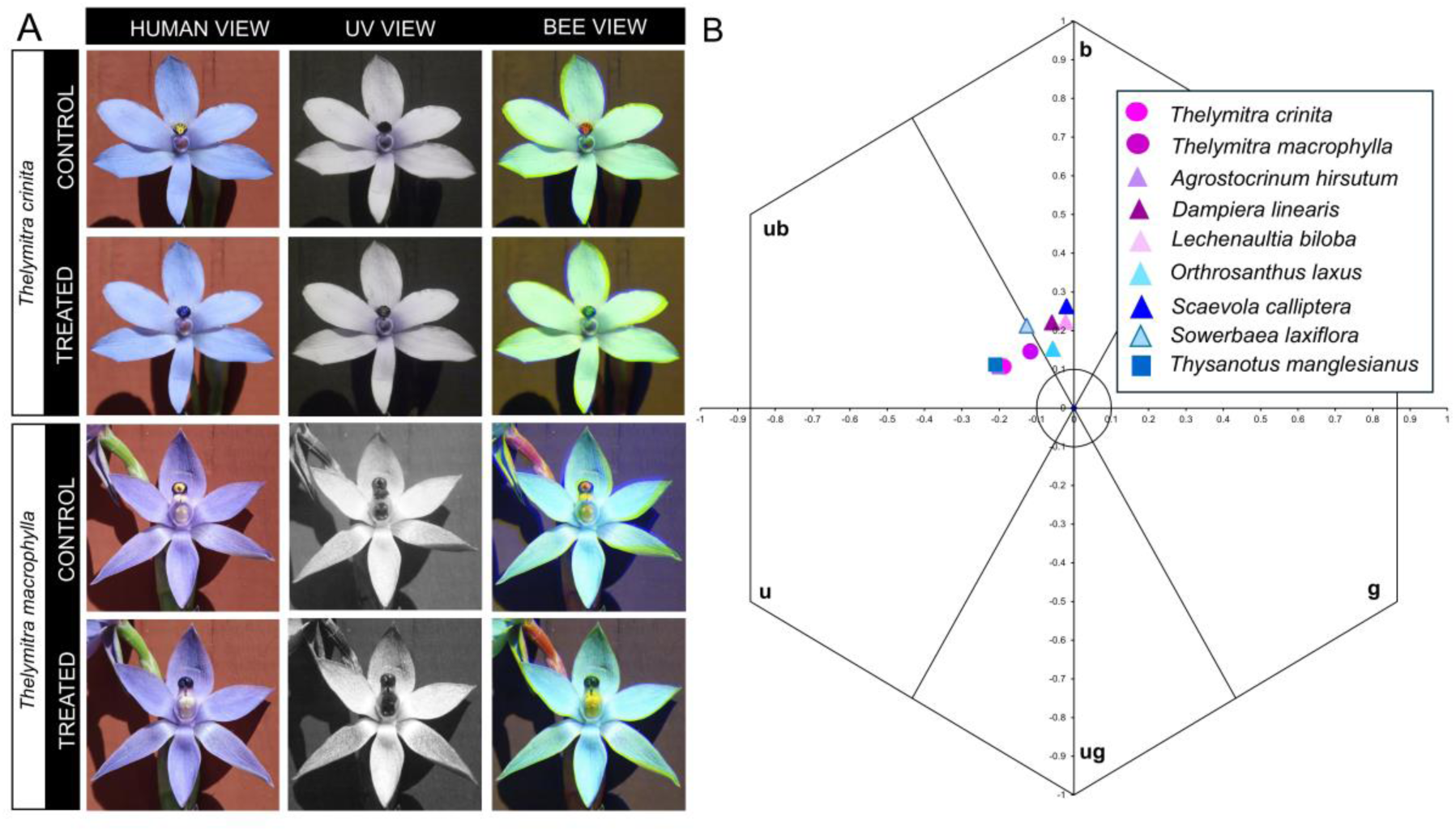
A: Natural (control) and painted (treated) flowers of *Thelymitra crinita* and *T. macrophylla* photographed in visible light (human view), UV, and transformed in false colour photography (bee view). B: Mean flower colour loci (n=6) of the deceptive orchids *T. crinita* and *T. macrophylla*, the co-flowering pollen-rewarding species (*Agrostocrinum hirsutum*, *Orthrosantus laxus, Sowerbaea laxiflora*, *Thysanotus manglesianus*) and the co-flowering nectar-rewarding species with similar flower colour (*Dampiera linearis*, *Lechenaultia biloba* and *Scaveola calliptera*). Calculations were based on the hexagon colour model of bee vision (u=ultraviolet, ub=ultraviolet-blue, b=blue, bg=blue-green; g=green; ug=ultraviolet-green; Chittka, 1992).

To test the ability for autogamous pollination in both orchids, we bagged the inflorescences on ten individuals of each species at the bud stage, and recorded fruit production four weeks later. None of the bagged inflorescences in any of the two species developed fruits.

### 2.6 Statistical analysis

We initially checked if number of flowers per plant differed between treatment groups (anther removal or paint vs. control) or populations at the onset of the experiments, by using two-way ANOVAs including the interaction between population and treatment. Number of flowers did not differ between treatments or populations in any of the experiments (all *P* > 0.31).

To examine the effects of treatment (anther removal or paint vs. control), population, and their interaction on number of fruits produced in the three experiments, we used Generalised Linear Models with Poisson distribution and log link. Data were in some cases zero-inflated, and we used a negative binomial distribution when we detected overdispersion of the residuals. We used the package glmm TMB in R Studio (version 4.2.3).

## 3 RESULTS

### 3.1 Flower colour

Analysis of spectral reflectance using the bee vision model showed that the average flower colour loci of *Thelymitra macrophylla* and *T. crinita* were located in the UV-blue region of the colour hexagon (Fig. 2B). All putative model species that were pollen-rewarding also grouped in this hexagon region, except for *Orthorosantus laxus,* which was located in the blue region. All nectar-rewarding species that co-flowered with *T. crinita* (*Dampiera linearis*, *Lechenaultia biloba* and *Scaveola calliptera*) also grouped in the blue region (Fig. 2B). The mean distances between colour loci of *T. macrophylla* and its pollen-rewarding putative model species were 0.06 (*S. laxiflora*), 0.06 (*O. laxus*), and 0.10 (*T. manglesianus*) hexagon units. The mean distances between colour loci of *T. crinita* and its pollen-rewarding model species were 0.01 (*A. hirsutum*) and 0.02 (*T. manglesianus*) units, and mean distances to the nectar-rewarding species were 0.17 (*D. linearis*), 0.20 (*L. biloba*), and 0.22 (*S. calliptera*; Table S3) units.

### 3.2 Pollinator observations

In the experimental arrays with *T. crinita*, a total of 118 insect approaches were observed (Table 1). The great majority of approaches were from bees, with more than half (n=34) by the genus *Leioproctus* (Colletidae). Individuals of *Leioproctus*, *Exoneura* (Apidae), and Halictidae landed on the orchid flowers, and attempted to hold and grasp the false anther with their mandibles and anterior legs, positioned with the head upwards or downwards (Fig. 1i). All observed bees carried pollen and were therefore females. Six of the bees manipulating the false anther (four *Leioproctus* and two *Exoneura* bees) appeared to attempt to buzz the orchid flower, but because visits lasted less than 2-3 seconds, we could ascertain buzzes for only two of the bees (based on emitted sound and observed wings vibrations). The observed bee behaviour on *T. crinita* was similar to the bee’s behaviour on flowers of the rewarding and buzz-pollinated species *T. manglesianus* and *A. hirsutum* (Video S1). We also observed two approaches to *T. crinita* by the blue-banded bee, *Amegilla cingulata* (Apidae), but we were unable to characterize their behaviour, due to very rapid visits (<1 second).

**Table 1.**
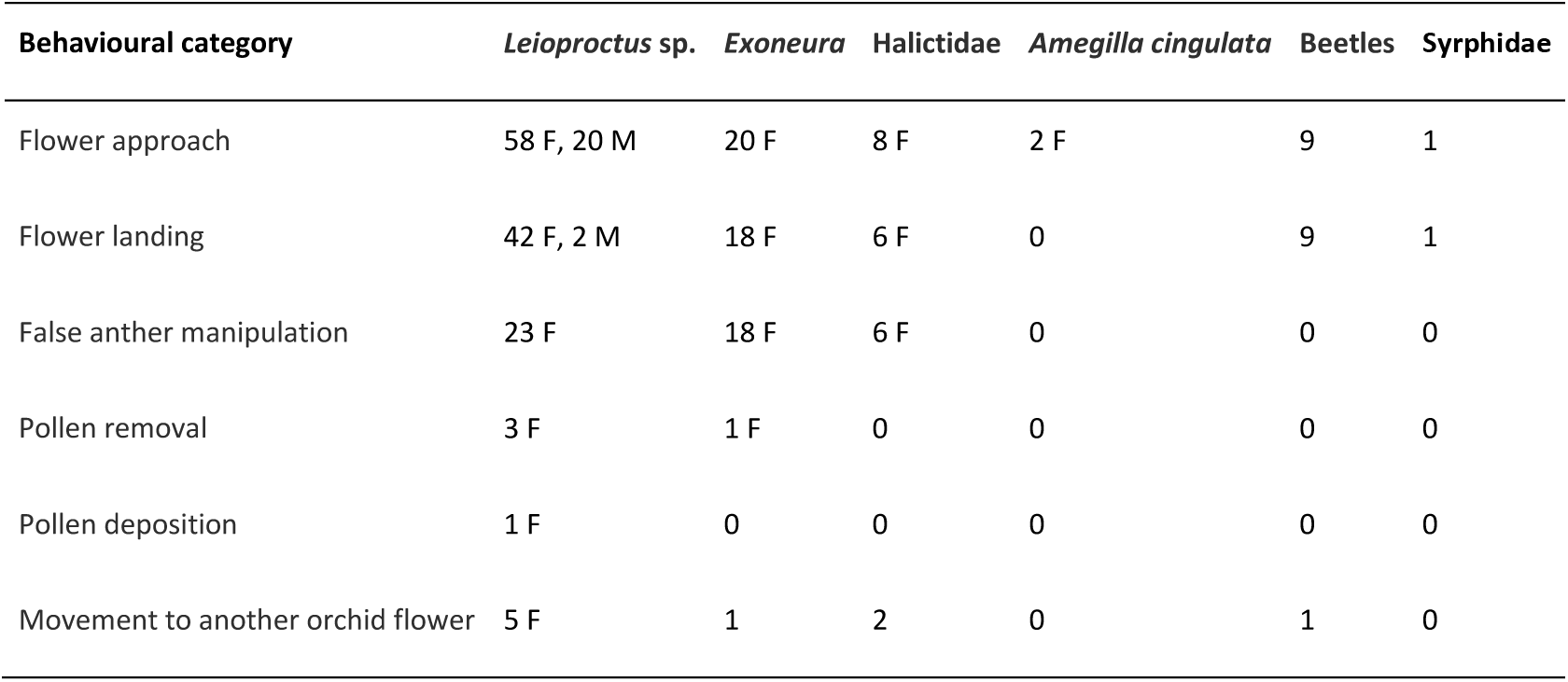
Number of bees, beetles, and hoverflies approaching and visting the orchid *Thelymitra crinita*. Behavioural categories reflect the sequential pollination process. M = male, F = female.

Only *Leioproctus* bees and one *Exoneura* bee were observed to remove and deposit orchid pollinia (Table 1). *Leioproctus* bees were positioned with the head upwards, and the tip of the abdomen touched the orchid reproductive structure (stigma and viscidium), resulting in pollinia attachment to the tip of the abdomen (Fig. S3). The *Exoneura* bee was observed removing orchid pollinia attached to the tip of its abdomen after spending six seconds over the false anther, moving its head up and down. None of the visiting bees carried any orchid pollinia on arrival to the flowers. Two of the *Leioproctus* bees that removed orchid pollinia moved directly to plants of *T. manglesianus*.

We observed a range of buzz-pollinating bees, including *Leioproctus*, *Exoneura*, and *A. cingulata*, visiting the pollen-rewarding putative model plants, *T. manglesianus* and *A. hirsutum*. A total of 62 and 54 bees were observed to produce wing vibrations and release pollen from anthers of *T. manglesianus* and *A. hirsutum*, respectively, and visits lasted up to 20 seconds (Video S2). All taxa observed to visit the orchids were also visiting the nectar-rewarding species in the study population, but none of the bees attempted to buzz these flowers.

### 3.3 False anther manipulations

a. *Anther removal experiment*

Fruit production of *T. macrophylla* was significantly reduced by the removal of false anthers, but was not affected by site, or the interaction between treatment and site (Table 2). In the site without model plants, mean fruit number was 66% lower in treated plants compared to controls, while in the site with models present, the corresponding reduction was 60% (Fig. 3A).

*Paint experiment*

**Figure 3.**
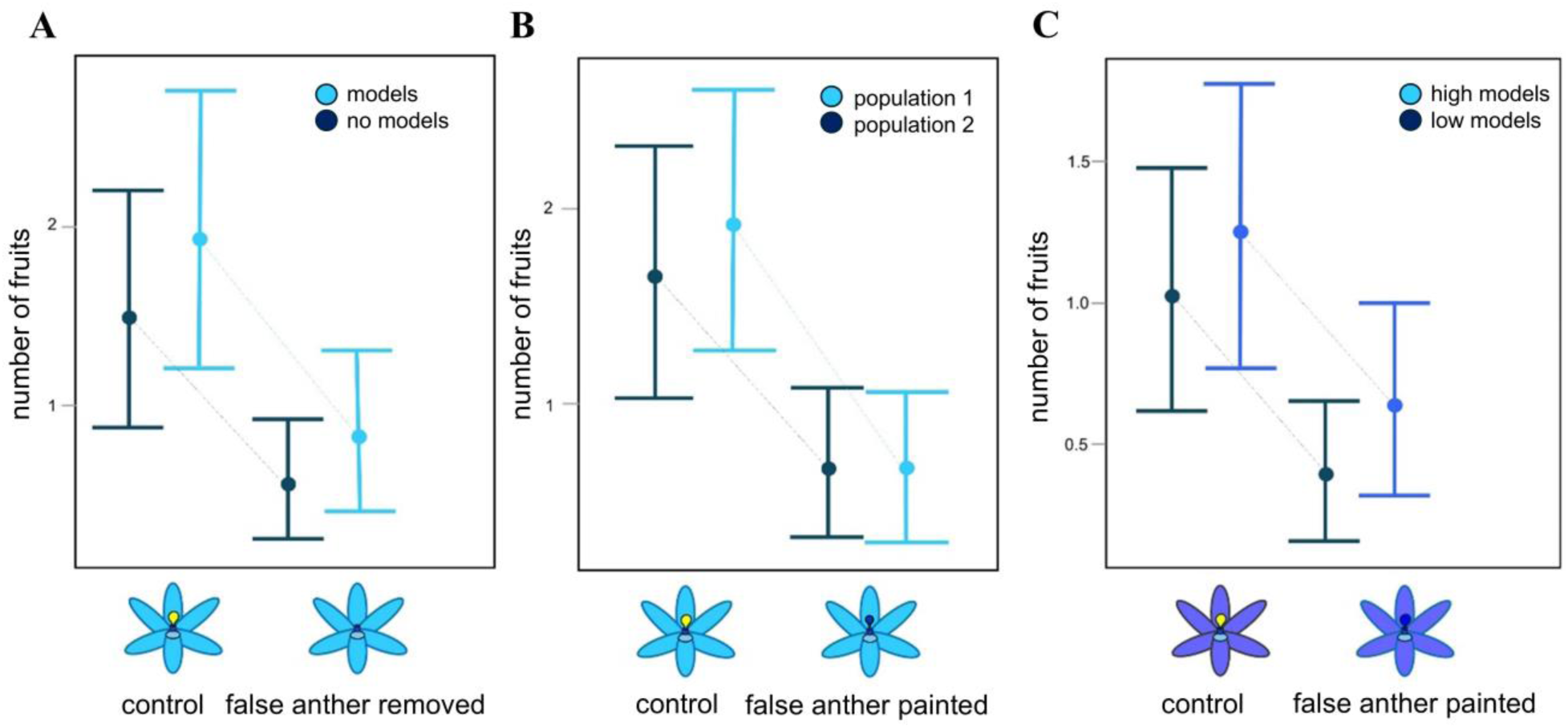
Effect of false anther manipulation (removal or paint) on number of fruits produced by individuals of *Thelymitra macrophylla* (panel A and B) and *T. crinita* (panel C). Symbols with confidential intervals indicate model estimates from generalized linear models with Poisson distribution and log link.

**Table 2.** The effect of population, treatment (anther removal or painting vs. control), and their interaction on fruit production of individuals of the orchids *Thelymitra macrophylla* (TM) and *T. crinita* (TC), analysed with generalized linear models with Poisson distribution and log link.

| Experiment | Effect | $\chi^2$ | p-value |
| --- | --- | --- | --- |
| Anther removal TM<br>n=94 | population | 1.64 | 0.20 |
|  | treatment | 13.92 | 0.0002 |
|  | pop*treatment | 0.08 | 0.78 |
| Anther paint TM<br>n=60 | population | 0.22 | 0.64 |
|  | treatment | 19.20 | <0.0001 |
|  | pop*treatment | 0.22 | 0.64 |
| Anther paint TC<br>n=96 | population | 1.88 | 0.17 |
|  | treatment | 11.65 | 0.0006 |
|  | pop*treatment | 0.43 | 0.51 |

Fruit production was significantly reduced by the painting of false anthers in both orchids, but was not affected by site, or the interaction between treatment and site (Table 2). In *T. macrophylla*, obscuring false anthers resulted in 63% and 71% reduction in number of fruits in the two sites that both had abundant model plants (Fig. 3B). In *T. crinita*, the painting treatment reduced fruit production by 66% in the site with low abundance of model plants, and by 51% in the site with high abundance of model plants (Fig. 3C).

## 4 DISCUSSION

We found that both sun orchids are visually similar to their co-flowering buzz-pollinated species, and observations of insect activity on *Thelymitra crinita* also suggest shared pollinators with buzz-pollinated species. In both sun orchid species, removing or obscuring the false anther caused strong reductions in female reproductive success. The results suggest that the false anther is a key pollination adaptation in the orchids and are consistent with stamen mimicry of buzz-pollinated plants in the rewarding community.

### 4.1 Adaptive value of false anthers

All three experiments involving anther manipulations showed that the false anther was a major determinant of female reproductive success of the orchid species. Fruit production in plants with excised or obscured anthers was on average reduced by 51-71%, indicating that this trait is key to visual attraction of pollinating bees. While bees are likely to be attracted to large floral displays from a distance (e.g., Benitez-Vieyra et al., 2006; Sletvold et al., 2010; Sletvold & Ågren, 2011), false anthers may be an important signal in proximity to the flowers. The colour contrast between the anther and petal could direct pollinators to the centre of the flower, and function as a ‘floral guide’ (Hansen et al., 2011; Lunau, 2017). Our insect observations revealed that most bees landed directly on the column containing the false anther, suggesting that bees mistake orchids for pollen-rewarding plants during their approach. It is also possible that the false anther influences bee behaviour after landing on the flower. Newman et al. (2022) found that anther mimics in *Tritonia laxifolia* (Iridaceae) not only contributed to pollinator attraction, but also influenced bee behaviour and the efficiency of pollen transfer. In sun orchids, the false anther likely induces the bee to bend its abdomen, ensuring contact between the tip of the abdomen and the orchid viscidium. We currently have too limited data on orchid pollinator visitation to evaluate the effect of the false anther on bee behaviour, but visiting bees are known to bend their abdomen on buzz-pollinated co-flowering *Thysanotus manglesianus* and *Agrostocrinum hirsutum*, resulting in pollen release (Eakin-Busher et al., 2017; Brudrett, 2024). This typical bee posture is associated with buzz pollination (Vallejo-Marín, 2019), and we expect that this posture is required to remove and deposit pollen in *Thelymitra* species, although further studies are necessary to confirm this. It has been suggested that flower-visiting bees do not exhibit pollen-collecting behaviour in response to stamen mimics (Vogel, 1978; Bernhardt, 1996; but see Wyatt & Sazima, 2011), but in this study, a few bees produced short buzzes on the sun orchid flowers. Bees are also known to attempt to collect pollen by buzzing pollen-depleted stamens of rewarding species (Burkart et al., 2014; Russell et al., 2017). Our limited data suggest that bees may buzz false anthers and given the potential prevalence of false anthers within *Thelymitra*, open opportunities for uncovering novel aspects of bee responses to this specific floral cue.

In line with theoretical predictions (Johnson & Schiestl, 2016) and previous empirical studies (Kunze & Gumbert, 2005; Anderson & Johnson, 2006), we expected the adaptive value of false anthers to be higher when putative model species were abundant. However, in both sun orchids, the fitness reduction following anther removal was similar across rewarding context. One possibility is that the difference in abundance of model plants was too limited to detect an effect. However, the anther removal experiment on *T. macrophylla* contrasted populations with and without model plants, and abundance of models differed by almost four times in the *T. crinita* experiment. Another possibility is that pollination success is affected by a combination of model and magnet effects, with the latter being dominant. For example, if nectar-producing magnet plants are absent, we might see low orchid reproductive success despite presence of pollen-rewarding model plants. Finally, it is possible that the spatial scale studied, i.e., the presence or abundance of putative model species in the local population, does not correspond to the relevant spatial scale influencing pollinator foraging decisions. In previous studies, focal scale has ranged from the local neighbourhood in the population (Anderson & Johnson, 2006) to the surrounding landscape (Duffy & Johnson, 2017; Scaccabarozzi et al., 2023), and studies manipulating rewarding context at several spatial scales would be a powerful approach to identify the most influential one. In addition, manipulations of the abundance of both putative model and magnet species could reveal their separate and interactive effects.

### 4.2 Mimicry of buzz-pollinated plants: a widespread phenomenon in Thelymitra?

The genus *Thelymitra* is characterized by star-shaped flowers with pseudoanthers, a morphology shared with many buzz-pollinated plants in the community. The colour analysis according to bee visual perception further showed that both orchids are visually similar to co-flowering buzz-pollinated flowers. The petal colour of *T. crinita* precisely matched the colour of *T. manglesianus* and *A. hirsutum,* with distances between colour loci of 0.01-0.02 hexagon units, considerably less than the 0.06 units required for bees to distinguish different colours (Dyer & Chittka 2004, Giurfa 2004). In contrast, none of the nectar-rewarding species that co-flowered with *T. crinita* had overlapping colour loci with the orchid (distance >0.1 units), suggesting that *T. crinita* specifically mimics buzz-pollinated plants. Also *T. macrophylla* matched the colour of its main co-flowering buzz-pollinated species, *S. laxiflora* and *O. laxus*, though not so closely as *T. crinita*. The average colour loci distance was 0.06 hexagon units to both species, which suggests that bees may be able to discriminate between models and mimic under certain conditions (Dyer & Chittka 2004). According to former studies*, T. macrophylla* is scented, with a fragrance similar to *O. laxus* (Edens-Meier et al., 2014). It is possible that in *T. macrophylla*, mimicry involves both visual and chemical cues, and that this combination of traits indicate a more generalised system (Johnson & Schiestl, 2016), compared to *T. crinita*.

Within Orchidaceae, *Thelymitra* exhibits characteristics that are uncommon within the family. Interestingly, some features are shared with the ancient genus *Apostasia*, which possess “solanum-type” flowers with an erect column bearing functional stamens (Kocyan & Endress, 2001; Zhang et al., 2017), suggested to be an adaptation to buzz pollination (Vogel, 1981; Kocyan & Endress, 2001). Shared traits include an undifferentiated labellum, star-like flowers, and fused anthers (which are non-functional in sun orchids). Similar features are also observed in other Asparagales (Zhang et al., 2017), including putative model plants of *Thelymitra* species. However, *Apostasia* is positioned at the base of the orchid clade (∼120–90 million years ago) (Chase, 2001; Givnish et al., 2016; Zhang et al., 2023; Escobar et al., 2024), while *Thelymitra* originated much later (∼10 million years ago) (Nauheimer et al., 2018). This large evolutionary distance indicates that the characteristic traits of sun orchids arose as novel adaptations to mimic buzz-pollinated model plants, rather than being due to shared ancestry. According to phylogenetic analysis by Nauheimer et al. (2018), *Thelymitra* and *Calochilus* are sister genera that originated from a common ancestor that also gave rise to their sister genus *Epiblema*. The latter shares the star-like floral morphology with *Thelymitra*, whereas *Calochilus* has bilateral flowers with a conspicuous “bearded” labellum. Neither *Epiblema* nor *Calochilus* displays the characteristic staminodal hood with its bright, contrasting tip (Burns-Balogh & Bernhardt, 1985), suggesting that this trait has evolved in *Thelymitra*. Further phylogenetic analysis of *Thelymitra* and closely related genera would be valuable to test whether the staminodal hood evolved once, followed by subsequent radiation, or multiple times independently. This question could also be explored by investigating whether shifts in pollination strategies align with changes in floral traits, with a particular focus on autogamous species within *Thelymitra*.

## 5 CONCLUSIONS

This study documented striking similarity between buzz-pollinated model plants and sun orchids in derived floral traits that significantly impact fitness. This supports the hypothesis of buzz-pollination mimicry. The strong reduction in orchid fitness when the false anther was removed was likely due to a reduction in attractiveness (absence of a floral guide signal) as well as in similarity to model plants, and to disentangle these effects, future experiments could reduce similarity to model plants without simultaneously diminishing attractiveness. Moreover, the lack of any effect of model abundance on orchid fitness shows that further studies are needed to determine how specific buzz-pollination mimicry is. While food-deceptive pollination strategies are common among orchids, *Thelymitra* appears to be unique in evolving a deceptive buzz-pollination strategy. The entire genus may be adapted to mimic buzz-pollinated flowers, a strategy particularly relevant in the Southwest Australian Floristic Region (SWAFR), where many bee genera (e.g., *Amegilla*, *Anthoglossa*, *Lasioglossum*, and *Leioproctus*) are capable of buzzing (Houston et al., 2018; Brundrett et al., 2024). With over 100 species and an established phylogeny, *Thelymitra* offers a fascinating opportunity to investigate the evolution, as well as the functional and adaptive significance of floral traits involved in mimicry.

## AUTHOR CONTRIBUTIONS

Conceptualization: Daniela Scaccabarozzi and Nina Sletvold. Investigation: Daniela Scaccabarozzi. Data curation: Daniela Scaccabarozzi and Nina Sletvold. Formal Analysis: Daniela Scaccabarozzi and Nina Sletvold. Project administration: Daniela Scaccabarozzi and Nina Sletvold. Supervision: Nina Sletvold. Writing-original draft Daniela Scaccabarozzi and Nina Sletvold.

## ACKNOWLEDGEMENTS

We thank Klaus Lunau for support with false colour photography and for providing comments on an earlier draft of the manuscript. We also thank Jon Ågren for his valuable comments on the manuscript and for revising the work. We thank Terry Houston at the Museum of Western Australia for insect identification, Mick Hurdus for providing photos of bees visiting orchids, and Max Massi and Peter Chapman for access to lab facilities and for lab assistance at Curtin University, School of Molecular and Life Sciences, Perth. We thank Annette Westhoff, Oma Fox, Graham Warren, and WANOSCG for advice on field sites, and the Department of Biodiversity, Conservation and Attractions for providing feedback on the research questions and proposed approach.

## STATEMENT OF INCLUSION

Our study brings together authors from different countries, including the lead scientist, who is partly based in the country where the study was carried out. All authors were authentically involved throughout the project. Whenever possible, our research was discussed with local stakeholders to seek feedback.

## REFERENCES

Anderson, B., & Johnson, S. D. (2006). The effects of floral mimics and models on each others’ fitness. Proceedings of the Royal Society B: Biological Sciences, 273(1589), 969–974. 10.1098/rspb.2005.3401

Beardsell, D. V., & Bernhardt, P. (1983). Pollination biology of Australian terrestrial orchids. In E. G Williams, R. B Knox, J. H. Gilbert & P. Bernhardt (Eds.), Pollination ’82 (pp. 166–183). University of Melbourne Press.

Bernhardt, P. & Burns-Balogh, P. (1986). Floral mimesis in *Thelymitra nuda* (Orchidaceae). Plant Systematics and evolution, 151, 187–202. 10.1007/BF02430274

Bernhardt, P. (1996). Anther adaptation in animal pollination. In W. G. D’Arcy & R. C. Keating (Eds), The anther form, function and phylogeny (pp. 192–220). Cambridge University Press.

Benitez-Vieyra, S., Medina, A. M., Glinos, E., & Cocucci, A. A. (2006). Pollinator-mediated selection on floral traits and size of floral display in *Cyclopogon elatus*, a sweat bee-pollinated orchid. Functional Ecology, 20(6), 948–957. 10.1111/j.1365-2435.2006.01179.x

Brundrett, M. C., Ladd, P. G., & Keighery, G. J. (2024). Pollination strategies are exceptionally complex in southwestern Australia–a globally significant ancient biodiversity hotspot. Australian Journal of Botany, 72(2). 10.1071/BT23007

Buchmann, S. L. (1983). Buzz pollination in angiosperms. Bo. Paper 266. https://digitalcommons.usu.edu/bee_lab_bo/266

Burkart, A., Schlindwein, C., & Lunau, K. (2014). Assessment of pollen reward and pollen availability in *Solanum stramoniifolium* and *Solanum paniculatum* for buzz-pollinating carpenter bees. Plant Biology, 16(2), 503–507. 10.1111/plb.12111

Chase M. 2001. The origin and biogeography of Orchidaceae. In A. M. Pridgeon, P. J. Cribb, M. W. Chase & F. N. Rasmussen (Eds.). Genera Orchidacearum: vol. 2. Orchidoideae (part one). Oxford University Press.

Chittka, L. (1992). The colour hexagon: a chromaticity diagram based on photoreceptor excitations as a generalized representation of colour opponency. Journal of Comparative Physiology A, 170, 533–543. 10.1007/BF00199331

Dafni, A., & Ivri, Y. (1981). The flower biology of *Cephalanthera longifolia* (Orchidaceae)—pollen imitation and facultative floral mimicry. Plant Systematics and Evolution, 137, 229–240. 10.1007/BF00982788

Dalziell, A. H., & Welbergen, J. A. (2016). Mimicry for all modalities. Ecology Letters, 19(6), 609–619. 10.1111/ele.12602

De Jager, M. L., & Anderson, B. (2019). When is resemblance mimicry?. Functional Ecology, 33(9), 1586–1596. 10.1111/1365-2435.13346

De Luca, P. A., & Vallejo-Marín, M. (2013). What’s the ‘buzz’ about? The ecology and evolutionary significance of buzz-pollination. Current opinion in plant biology, 16(4), 429–435. 10.1016/j.pbi.2013.05.002

Dieringer, G., & Cabrera R, L. (2002). The interaction between pollinator size and the bristle staminode of Penstemon digitalis (Scrophulariaceae). American Journal of Botany, 89(6), 991–997. 10.3732/ajb.89.6.991

Duffy, K. J., & Johnson, S. D. (2017). Effects of distance from models on the fitness of floral mimics. Plant Biology, 19(3), 438–443. 10.1111/plb.12555

Dyer, A. G., & Chittka, L. (2004). Biological significance of distinguishing between similar colours in spectrally variable illumination: bumblebees (*Bombus terrestris*) as a case study. Journal of Comparative Physiology A, 90(2), 105–14. 10.1007/s00359-003-0475-2

Duncan, D. H., Cunningham, S. A., & Nicotra, A. B. (2004). High self-pollen transfer and low fruit set in buzz-pollinated *Dianella revoluta* (Phormiaceae). Australian Journal of Botany, 52(2), 185–193. 10.1071/BT03139

Edens-Meier, R., Westhus, E., & Bernhardt, P. (2013). Floral biology of large-flowered *Thelymitra* species (Orchidaceae) and their hybrids in Western Australia. Telopea, 15. 10.7751/telopea2013020

Edens-Meier, R. M., Raguso, R. A., Westhus, E., & Bernhardt, P. (2014). Floral fraudulence: do blue *Thelymitra* species (Orchidaceae) mimic *Orthrosanthus laxus* (Iridaceae)?. Telopea, 17, 15–28. 10.7751/telopea20147392

Eakin-Busher, E. L., Fontaine, J. B., & Ladd, P. G. (2016). The bees don’t know and the flowers don’t care: the effect of heterospecific pollen on reproduction in co-occurring *Thysanotus* species (Asparagaceae) with similar flowers. Botanical Journal of the Linnean Society, 181(4), 640–650. 10.1111/boj.12433

Feinsinger, P. (1987). Effects of plant species on each other’s pollination: is community structure influenced?. Trends in Ecology & Evolution, 2(5), 123–126. 10.1016/0169-5347(87)90052-8

Givnish, T. J., Spalink, D., Ames, M., Lyon, S. P., Hunter, S. J., Zuluaga, A., Doucette, A., Caro, G. G., McDaniel, J., Clements, M. A., Arroyo, M. T. K., Endara, L., Kriebel, R., Williams, N. H., & Cameron, K. M. (2016). Orchid historical biogeography, diversification, Antarctica and the paradox of orchid dispersal. Journal of Biogeography, 43(10), 1905–1916. 10.1111/jbi.12854

Giurfa, M. (2004). Conditioning procedure and color discrimination in the honeybee *Apis mellifera*. Naturwissenschaften, 91, 228–231. 10.1007/s00114-004-0530-z

Hopper, S. D., & Gioia, P. (2004). The southwest Australian floristic region: evolution and conservation of a global hot spot of biodiversity. The Annual Review of Ecology, Evolution, and Systematics, 35, 623–650. 10.1146/annurev.ecolsys.35.112202.130201

Houston, T. (2018). A guide to native bees of Australia. Csiro Publishing.

Hansen, D. M., Van der Niet, T., & Johnson, S. D. (2012). Floral signposts: testing the significance of visual ‘nectar guides’ for pollinator behaviour and plant fitness. Proceedings of the Royal Society B: Biological Sciences, 279(1729), 634–639. 10.1098/rspb.2011.1349

Jamie, G. A. (2017). Signals, cues and the nature of mimicry. Proceedings of the Royal Society B: Biological Sciences, 284(1849), 20162080. 10.1098/rspb.2016.2080

Jersáková, J., Johnson, S.D., & Jürgens, A. (2009). Deceptive Behavior in Plants. II. Food Deception by Plants: From Generalized Systems to Specialized Floral Mimicry. In F. Balka, (Eds.), Plant-Environment Interactions. Signaling and Communication in Plants. Springer. https://doi.org/10.1007/978-3-540-89230-4_12

Jersáková, J., Jürgens, A., Šmilauer, P., & Johnson, S. D. (2012). The evolution of floral mimicry: identifying traits that visually attract pollinators. Functional Ecology, 26(6), 1381–1389. 10.1111/j.1365-2435.2012.02059.x

Jersáková, J., Spaethe, J., Streinzer, M., Neumayer, J., Paulus, H., Dötterl, S., & Johnson, S. D. (2016). Does *Traunsteinera globosa* (the globe orchid) dupe its pollinators through generalized food deception or mimicry?. Botanical journal of the Linnean Society, 180(2), 269–294. 10.1111/boj.12364

Johnson, S. D. (1994). Evidence for Batesian mimicry in a butterfly-pollinated orchid. Biological journal of the Linnean Society, 53(1), 91–104. 10.1111/j.1095-8312.1994.tb01003.x

Johnson, S. D. (2000). Batesian mimicry in the non-rewarding orchid Disa pulchra, and its consequences for pollinator behaviour. Biological Journal of the Linnean Society, 71(1), 119–132. 10.1111/j.1095-8312.2000.tb01246.x

Johnson, S. D., Peter, C. I., Nilsson, L. A., & Ågren, J.(2003a). Pollination success in a deceptive orchid is enhanced by co-occurring rewarding magnet plants. Ecology 84(11), 2919–2927.

Johnson, S. D., Alexandersson, R., & Linder, H. P. (2003b). Experimental and phylogenetic evidence for floral mimicry in a guild of fly-pollinated plants. Biological Journal of the Linnean Society, 80(2), 289–304. 10.1046/j.1095-8312.2003.00236.x

Johnson, S. D., Peter, C. I., Nilsson, L. A., & Ågren, J. (2003). Pollination success in a deceptive orchid is enhanced by co-occurring rewarding magnet plants. Ecology, 84(11), 2919–2927. 10.1890/02-0471

Johnson, S. D. & Schiestl, F. P. (2016). Floral mimicry. Oxford University Press.

Joron, M., & Mallet, J. L. (1998). Diversity in mimicry: paradox or paradigm?. Trends in ecology & evolution, 13(11), 461–466. 10.1016/S0169-5347(98)01483-9

Kearns, C. A., & Inouye, D. W. (1993). Techniques for pollination biologists. University Press of Colorado.

Kocyan, A., & Endress, P. K. (2001). Floral structure and development of *Apostasia* and *Neuwiedia* (Apostasioideae) and their relationships to other Orchidaceae. International Journal of Plant Sciences, 162(5), 847–867. 10.1086/320781

Kunze, J., & Gumbert, A. (2001). The combined effect of color and odor on flower choice behavior of bumble bees in flower mimicry systems. Behavioral Ecology, 12(4), 447–456. 10.1093/beheco/12.4.447

Laverty, T. M. (1992). Plant interactions for pollinator visits: a test of the magnet species effect. Oecologia, 89, 502–508. 10.1007/BF00317156

Lunau, K., & Wester, P. (2017). Mimicry and deception in pollination. In G. Becard G (Eds.), Advances in Botanical Research, Vol. 82, How Plants Communicate with their Biotic Environment (pp. 259–279). Academic Press.

Lunau, K. (2000). The ecology and evolution of visual pollen signals. Plant Systematics and Evolution, 222, 89–111. 10.1007/BF00984097

Lunau, K., Konzmann, S., Winter, L., Kamphausen, V., & Ren, Z. X. (2017). Pollen and stamen mimicry: the alpine flora as a case study. Arthropod-Plant Interactions, 11, 427–447. 10.1007/s11829-017-9525-5

Lunau, K., Scaccabarozzi, D., Willing, L., & Dixon, K. (2021). A bee’s eye view of remarkable floral colour patterns in the south-west Australian biodiversity hotspot revealed by false colour photography. Annals of Botany, 128(7), 821–824. 10.1093/aob/mcab088

Lunau, K., Ren, Z. X., Fan, X. Q., Trunschke, J., Pyke, G. H., & Wang, H. (2020). Nectar mimicry: a new phenomenon. Scientific Reports, 10(1), 7039. 10.1038/s41598-020-63997-3

Lunau K, Brito VLG, Camargo MGG (2024) Pollen, anther, stamen, and androecium mimicry. Plant Biology 26: 349–368. 10.1111/plb.13628

Myers, N., Mittermeier, R. A., Mittermeier, C. G., Da Fonseca, G. A., & Kent, J. (2000). Biodiversity hotspots for conservation priorities. Nature, 403(6772), 853–858. 10.1038/35002501

Newman, E. L., Khoury, K. L., Niekerk, S. E. V., & Peter, C. I. (2022). Structural anther mimics improve reproductive success through dishonest signalling that enhances both attraction and the morphological fit of pollinators with flowers. Evolution, 76(8), 1749–1761. 10.1111/evo.14540

Pérez-Escobar, O. A., Bogarín, D., Przelomska, N. A., Ackerman, J. D., Balbuena, J. A., Bellot, S., … & Antonelli, A. (2024). The origin and speciation of orchids. New Phytologist, 242(2), 700–716. 10.1111/nph.19580

Peter, C. I., & Johnson, S. D. (2009). Reproductive biology of *Acrolophia cochlearis* (Orchidaceae): estimating rates of cross-pollination in epidendroid orchids. Annals of Botany, 104(3), 573–581. 10.1093/aob/mcn218

Pridgeon, A. M., Cribb, P. J., Chase, M. W., & Rasmussen, F. N. (2001). Genera ’Orchidacearum’. Vol. 2. Oxford University Press.

Renner, S.S., 2006. Rewardless flowers in the angiosperms and the role of insect cognition in their evolution. Plant-pollinator interactions: from specialization to generalization (pp.123–144). University of Chicago Press.

Rakosy, D., Cuervo, M., Paulus, H. F., & Ayasse, M. (2017). Looks matter: changes in flower form affect pollination effectiveness in a sexually deceptive orchid. Journal of Evolutionary Biology, 30(11), 1978–1993. 10.1111/jeb.13153

Roy, B. A., & Widmer, A. (1999). Floral mimicry: a fascinating yet poorly understood phenomenon. Trends in Plant Science, 4(8), 325–330. 10.1016/S1360-1385(99)01445-4

Russell, A. L., Buchmann, S. L., & Papaj, D. R. (2017). How a generalist bee achieves high efficiency of pollen collection on diverse floral resources. Behavioral Ecology, 28(4), 991–1003. 10.1093/beheco/arx058

Ruxton, G. D., & Schaefer, H. M. (2011). Alternative explanations for apparent mimicry. Journal of ecology, 99(4), 899–904. 10.1111/j.1365-2745.2011.01806.x

Scaccabarozzi, D., Cozzolino, S., Guzzetti, L., Galimberti, A., Milne, L., Dixon, K. W., & Phillips, R. D. (2018). Masquerading as pea plants: behavioural and morphological evidence for mimicry of multiple models in an Australian orchid. Annals of botany, 122(6), 1061–1073. 10.1093/aob/mcy166

Scaccabarozzi, D., Guzzetti, L., Phillips, R. D., Milne, L., Tommasi, N., Cozzolino, S., & Dixon, K. W. (2020a). Ecological factors driving pollination success in an orchid that mimics a range of Fabaceae. Botanical Journal of the Linnean Society, 194(2), 253–269. 10.1093/botlinnean/boaa039

Scaccabarozzi, D., Dixon, K. W., Tomlinson, S., Milne, L., Bohman, B., Phillips, R. D., & Cozzolino, S. (2020b). Pronounced differences in visitation by potential pollinators to co-occurring species of Fabaceae in the Southwest Australian biodiversity hotspot. Botanical Journal of the Linnean Society, 194(3), 308–325. 10.1093/botlinnean/boaa053

Scaccabarozzi, D., Galimberti, A., Dixon, K. W., & Cozzolino, S. (2020c). Rotating Arrays of Orchid Flowers: A simple and effective method for studying pollination in food deceptive plants. Diversity, 12(8), 286. 10.3390/d12080286

Scaccabarozzi, D., Lunau, K., Guzzetti, L., Cozzolino, S., Dyer, A. G., Tommasi, N., Biella, P., Galimberti, A., Labra, M., Bruni, I., Pattarini, G., Brundrett, M., & Gagliano, M. (2023). Mimicking orchids lure bees from afar with exaggerated ultraviolet signals. Ecology and evolution, 13(1), e9759. 10.1002/ece3.9759

Scaccabarozzi, D., Guzzetti, L., Pioltelli, E., Brundrett, M., Aromatisi, A., Polverino, G., Vallejo-Marín, M., Cozzolino, S., & Ren, Z. X. (2024). Evidence of introduced honeybees (*Apis mellifera*) as pollen wasters in orchid pollination. Scientific Reports, 14(1), 14076. 10.1038/s41598-024-64218-x

Sletvold, N., Grindeland, J. M., & Ågren, J. (2010). Pollinator-mediated selection on floral display, spur length and flowering phenology in the deceptive orchid *Dactylorhiza lapponica*. New Phytologist, 188(2), 385–392. 10.1111/j.1469-8137.2010.03296.x

Sletvold, N., & Ågren, J. (2011). Nonadditive effects of floral display and spur length on reproductive success in a deceptive orchid. Ecology, 92(12), 2167–2174. 10.1890/11-0791.1

Thomson, J. D. (1978). Effects of stand composition on insect visitation in two-species mixtures of *Hieracium*. American Midland Naturalist, 100(2) 431–440. 10.2307/2424843

Vallejo-Marín, M. (2019). Buzz pollination: studying bee vibrations on flowers. New Phytologist, 224(3), 1068–1074. 10.1111/nph.15666

Verhoeven, C., Ren, Z. X., & Lunau, K. (2018). False-colour photography: a novel digital approach to visualize the bee view of flowers. Journal of Pollination Ecology, 23, 102–118. 10.26786/1920-7603(2018)11

Vogel, S. (1981). Bestäubungskonzepte der Monokotylen und ihr Ausdruck im System. Berichte der Deutschen Botanischen Gesellschaft, 94(1), 663–675.

Walker-Larsen, J., & Harder, L. D. (2000). The evolution of staminodes in angiosperms: patterns of stamen reduction, loss, and functional re-invention. American Journal of Botany, 87(10), 1367–1384. 10.1111/j.1438-8677.1981.tb03434.x

Wen, S-J., Chen, S., Rech, A. R., Wang, H., Wang, Z., Wu, D., & Ren, Z-X. (2024). The functional dilemma of nectar mimic staminodes in *Parnassia wightiana* (Celastraceae): Attracting pollinators and florivorous beetles. Ecology and Evolution, 14:e70380. 10.1002/ece3.70380

Wyatt, G. E., & Sazima, M. (2011). Pollination and reproductive biology of thirteen species of Begonia in the Serra do Mar State Park, São Paulo, Brazil. Journal of Pollination Ecology, 6. 10.26786/1920-7603(2011)16

Western Australian Herbarium (1998). Florabase, The Western Australian Flora, Department of Biodiversity, Conservation and Attractions. https://florabase.dbca.wa.gov.au/ (Accessed 14 September 2024).

Zhang, G. Q., Liu, K. W., Li, Z., Lohaus, R., Hsiao, Y. Y., Niu, S. C., … & Liu, Z. J.. (2017). The Apostasia genome and the evolution of orchids. Nature, 549(7672), 379–383. 10.1038/nature23897

