## Supplementary material for "Do sun orchids mimic buzz-pollinated plants? An experimental test of the adaptive significance of false anthers"

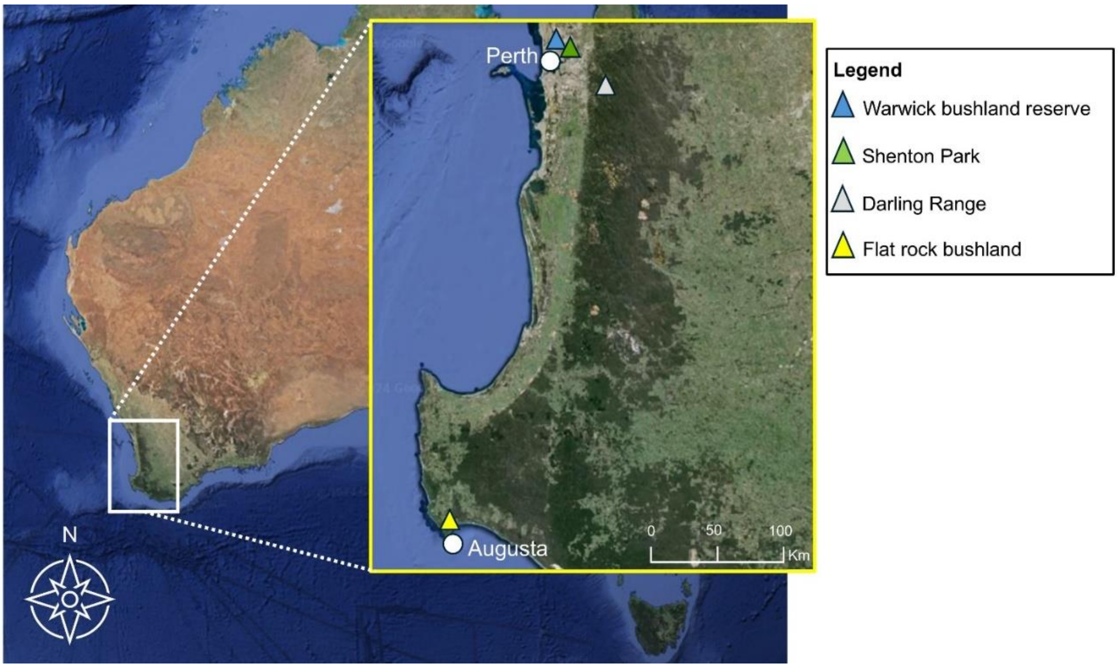


**Figure S1.** The geographical location of the *Thelymitra* study populations in Southwestern Australia. Populations of *T. macrophylla* are located in Warwick bushland Reserve (blue symbol) and Shenton Park (green symbol) in Perth*,* while study population of *T. crinita* are located in the Darling Range of Perth (grey symbol) and in Flat rock bushland in Augusta (yellow symbol).

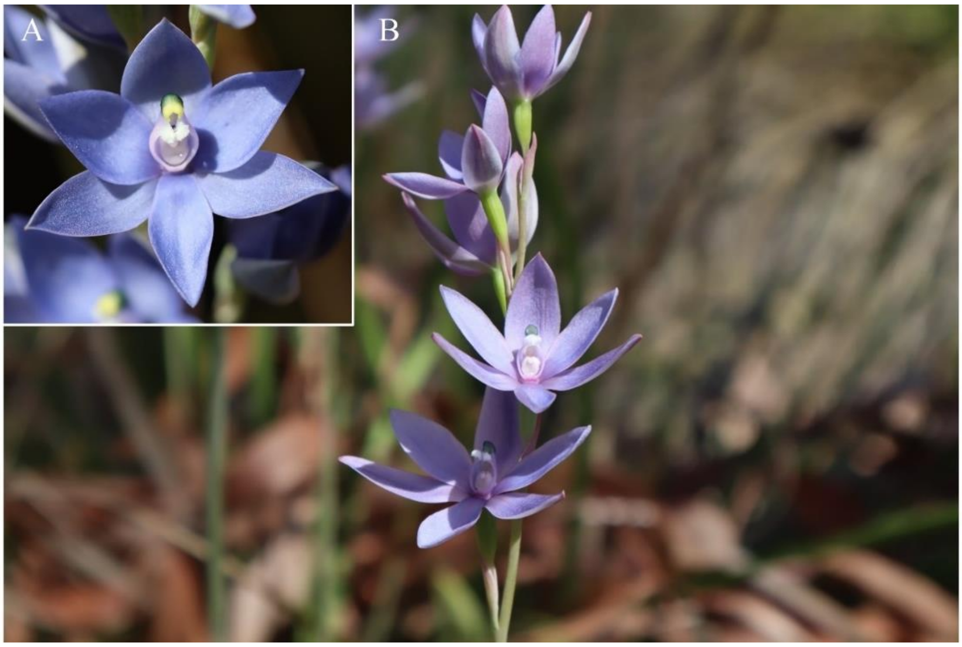


**Figure S2.** An illustration of the intact false anther (inlet A) and anther removal treatment in *Thelymitra macrophylla* plants, where false anthers (the upper part of the column) were excised from all open flowers using scissors (photo B).

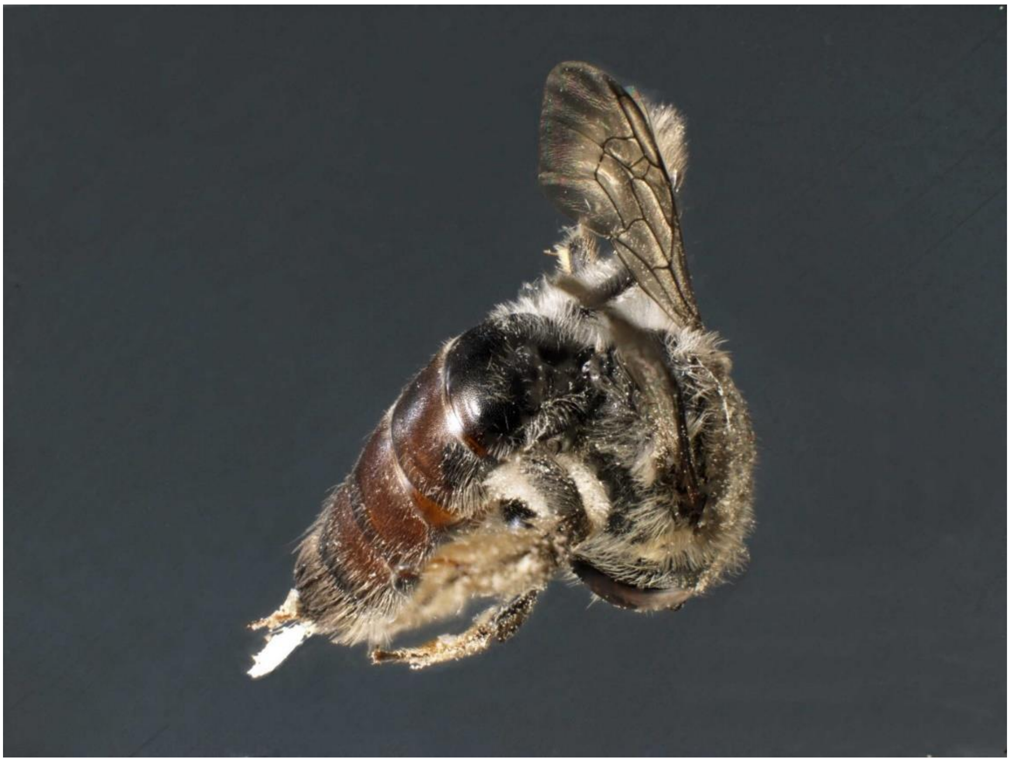

**Figure S3.** Female bee of *Leioproctus* (Subgenus: Odontocollete; Colletidae) carrying pollinia of *Thelymitra crinita*. Body length: 10 mm. Specimen identified and deposited at WA Museum, Code TC02.

**Table S1.** Mean and standard deviation of colour loci coordinates for each included plant species (n= 6 individuals per species): the sun orchids *Thelymitra crinita* and *T. macrophylla,* their coflowering pollen-rewarding plants (*Agrostocrinum hirsutum, Orthrosantus laxus, Sowerbaea laxiflora, Thysanotus manglesianus*), and other coflowering nectar-rewarding species with similar flower colour (*Dampiera linearis, Lechenaultia biloba, Scaveola calliptera*). Colour loci were calculated using the hexagon colour model of bee vision (Chittka, 1992).

| **Species** | **Mean (x)** | **Mean (y)** | **SD (x)** | **SD (y)** |
|---|---|---|---|---|
| *Thelymitra crinita* | -0.19 | 0.11 | 0.02 | 0.01 |
| *Thelymitra macrophylla* | -0.12 | 0.15 | 0.01 | 0.01 |
| *Agrostocrinum hirsutum* | -0.20 | 0.10 | 0.02 | 0.03 |
| *Dampiera linearis* | -0.06 | 0.22 | 0.03 | 0.02 |
| *Lechenaultia biloba* | -0.02 | 0.22 | 0.02 | 0.01 |
| *Thysanotus manglesianus* | -0.21 | 0.11 | 0.01 | 0.01 |
| *Scaveola calliptera* | -0.02 | 0.26 | 0.01 | 0.01 |
| *Orthorosanthis laxus* | -0.06 | 0.15 | 0.09 | 0.06 |
| *Sowerbea laxiflora* | -0.12 | 0.21 | 0.07 | 0.02 |

**Table S2.** Insects caught on *Thelymitra crinita*, *T. macrophylla, Thysanotus manglesianus* and *Agrostocrinum hirsutum.* All specimens were sexed and identified at the Western Australia Museum, at genus level. In the 'Sex' column, 'F' indicates female, 'M' indicates male, and 'na' indicates not available. *Orchid pollinia on the bee abdomen.

| Code | Date | Site | Plant species | Insect taxon | Sex |
|---|---|---|---|---|---|
| AG 01 | 1/10/2023 | Canning Rd - Kalamunda | *Agrosocrinum hirsutum* | *Leioproctus* | F |
| TS 02 | 29/09/2023 | Canning Rd - Kalamunda | *Thysonatus manglesianus* | *Exoneura* | F |
| TS 03 | 29/09/2023 | Canning Rd - Kalamunda | *Thysonatus manglesianus* | *Exoneura* | F |
| TM 01 | 22/09/2023 | Shantonk Park | *Thelymitra macrophylla* | *Liparetrus* | na |
| TM 02 | 22/09/2023 | Shantonk Park | *Thelymitra macrophylla* | *Liparetrus* | na |
| TM 03 | 22/09/2023 | Shantonk Park | *Thelymitra macrophylla* | *Liparetrus* | na |
| TM 04 | 22/09/2023 | Shantonk Park | *Thelymitra macrophylla* | *Liparetrus* | na |
| TM 05 | 22/09/2023 | Shantonk Park | *Thelymitra macrophylla* | *Liparetrus* | na |
| TM 06 | 22/09/2023 | Shantonk Park | *Thelymitra macrophylla* | *Liparetrus* | na |
| TC 01 | 2/11/2023 | Augusta flat rock | *Thelymitra crinita* | *Leioproctus* (Odontocolletes) | M |
| TC 02 | 3/11/2023 | Augusta flat rock | *Thelymitra crinita* | *Leioproctus* (Odontocolletes)* | F |
| TC 03 | 3/11/2023 | Augusta flat rock | *Thelymitra crinita* | *Leioproctus* (Odontocolletes)* | F |
| TC 04 | 21/10/2024 | Canning Rd - Kalamunda | *Thelymitra crinita* | *Exoneura* | F |
| TC 05 | 28/10/2024 | Margaret river | *Thelymitra crinita* | *Exoneura** | F |
| TS 04 | 3/11/2023 | Augusta flat rock | *Thysonatus manglesianus* | *Leioproctus* (Odontocolletes) | F |
| TS 05 | 3/11/2023 | Augusta flat rock | *Thysonatus manglesianus* | *Leioproctus* (Odontocolletes) | F |

### Supplementary results

**Video S1.** Bees of genera *Leioproctus* (Colletidae) patrolling (=males inspecting flowers), landing, and manipulating false anthers, or attempting to buzz on flowers of *Thelymitra crinita*. Bees of genus *Exoneura* (Colletidae) and Halictidae family manipulating false anthers or attempting to buzz on flowers of *T. crinita*.
https://youtu.be/SZr2r4GwgdE

**Video S2.** Bees of genera *Leioproctus* (Colletidae) and *Exoneura* (Apidae) visiting and buzzing on flowers of model plants, *Thysonatus manglesianus* and *Agrosocrinum hirsutum.*
https://youtu.be/V5bJ7PXF1TU
